# High-throughput analysis of B3GLCT regulation predicts phenotype of Peters’ Plus Syndrome in line with the miRNA Proxy Hypothesis

**DOI:** 10.1101/2021.04.01.438139

**Authors:** Chu T. Thu, Jonathan Y. Chung, Deepika Dhawan, Christopher A. Vaiana, Lara K. Mahal

## Abstract

MicroRNAs (miRNAs, miRs) finely tune protein expression and target networks of 100s-1000s of genes that control specific biological processes. They are critical regulators of glycosylation, one of the most diverse and abundant posttranslational modifications. In recent work, miRs have been shown to predict the biological functions of glycosylation enzymes, leading to the “miRNA proxy hypothesis” which states, “if a miR drives a specific biological phenotype…, the targets of that miR will drive the same biological phenotype.” Testing of this powerful hypothesis is hampered by our lack of knowledge about miR targets. Target prediction suffers from low accuracy and a high false prediction rate. Herein, we develop a high-throughput experimental platform to analyze miR:target interactions, miRFluR. We utilize this system to analyze the interactions of the entire human miRome with beta-3-glucosyltransferase (B3GLCT), a glycosylation enzyme whose loss underpins the congenital disorder Peters’ Plus Syndrome. Although this enzyme is predicted by multiple algorithms to be highly targeted by miRs, we identify only 27 miRs that downregulate B3GLCT, a >96% false positive rate for prediction. Functional enrichment analysis of these validated miRs predict phenotypes associated with Peters’ Plus Syndrome, although B3GLCT is not in their known target network. Thus, biological phenotypes driven by B3GLCT may be driven by the target networks of miRs that regulate this enzyme, providing additional evidence for the miRNA Proxy Hypothesis.

## INTRODUCTION

MicroRNAs (miRNAs, miRs) are small non-coding RNAs which fine-tune protein expression through binding messenger RNA (mRNA), primarily via the 3’-untranslated region (3’-UTR), downregulating protein levels. They are thought to target networks of 100s-1000s of genes that control specific biological processes, tightening their expression range and dampening noisy expression ^1, 2^. Currently, ~2700 human miRs have been reported, although the functions and targets of many are unknown. Contemporary knowledge of miR:target interactions relies heavily on computational tools and prediction algorithms such as Targetscan ^3^ and miRwalk ^4^ to identify interaction pairs. Only ~ 0.01% of all predicted human miR:target interactions have been validated to date ^5^. Limited studies comparing prediction to experimental data has estimated that between 16-63% of all predictions identify functional miR:target interactions ^6, 7^. However, these studies focused mainly on a limited set of well-known cancer genes and a subset of abundant and well-studied miRs, which may bias the analysis. To date, no study has comprehensively analyzed the human miRome regulation of a gene.

miRs are emerging as critical regulators of the glycome ^8–11^. Glycosylation is one of the most abundant and diverse post-translational modifications with roles in almost every disease state ^12^. However, identifying which glycosylation enzyme underlies which glycan epitope and concordant biology is still a barrier to our understanding of the glycome. Previous work from our laboratory identified miR regulatory networks that control glycosylation, arguing miRs are major regulators of the glycome ^9^. Downregulation of miR targets commonly recapitulates the phenotype induced by a miR. We realized that this might enable us to identify biological phenotypes of specific glycogenes, a point we demonstrated in work by Kurcon et al. ^10^. In this paper, we showed that glycosylation enzymes targeted by the miR-200 family, which are known to impact epithelia to mesenchymal transition (EMT) and migration, also regulate EMT and migration. This led us to formulate the “miRNA proxy hypothesis” which states, “if a miR drives a specific biological phenotype., the targets of that miR will drive the same biological phenotype. Thus, miRs can be used to identify (by proxy) the biological functions of specific glycosylation enzymes (or other proteins).” ^8^ We used this approach to identify glycosylation enzymes controlling cell cycle, providing additional evidence for our hypothesis ^11^. Testing of this hypothesis and utilization of this approach to identify the biological functions of glycosylation enzymes requires a thorough knowledge of miR:target interactions. However, in our original work, we found that only 3 out of the 11 miR:target interactions identified by prediction were accurate, and discovered 4 unpredicted interactions ^9^. The high false positive rates of prediction observed, coupled with significant false negatives, points to the need for more accurate data on miR regulation of glycosylation enzymes before we can test and utilize our hypothesis.

To overcome these obstacles, we developed a system for the experimental mapping of miR:target interactions, the miR-FluR high-throughput assay. This assay utilizes a genetically encoded **<u>flu</u>**orescent **<u>r</u>**atiometric reporter to identify **miR**: 3’-UTR interactions in a 384-well plate assay, enabling interrogation of the entire human miRome. We applied this assay to mapping the miR regulation of beta-3-glucosyltranferase (B3GLCT). This enzyme is predicted to be highly regulated. However, we show that B3GLCT predictions have a high false positive rate in multiple algorithms (>96%), implying that prediction accuracy may be far worse than currently thought. Previously, we demonstrated that this enzyme is regulated by the miR-200 family and has a role in EMT ^10^. This was consistent with the cleft palate observed in Peters’ Plus Syndrome, a genetic disorder caused by loss of B3GLCT. Herein, we find that the set of miRs downregulating B3GLCT predict multiple aspects of Peter’s Plus Syndrome, in line with our miRNA Proxy Hypothesis.

## RESULTS AND DISCUSSION

### Development of miRFluR high-throughput assay

To date, high-throughput analyses of miR-mRNA interactions have focused on the use of either luciferase assays in a 96-well format ^6^ or HITS-CLIP analysis, which pulls down miRs associated with mRNA in RISC complexes ^14^. Luciferase assays require the lysis of cells and expensive reagents. In contrast, HITS-CLIP assays focus on a specific miR, are cell-type dependent, and do not readily identify miR interactions with low abundance genes ^10, 15^. To overcome these limitations, we developed a **<u>fluo</u>**rescent **<u>r</u>**atiometric assay to identify **miR**: 3’-UTR interactions using a genetically encoded fluorescent reporter (miRFluR). Although both single and dual-color genetically encoded fluorescent reporters have been used to study miRs in live cells, their use has been limited to examining single miR:mRNA interactions by microscopy or flow cytometry ^16, 17^.

We created an optimized dual-color reporter, pFmiR, for high-throughput analysis of the regulatory interactions of the human miRome (pFmiR-3’UTR, **Figure 1A and S1**). In pFmiR, the 3’UTR of a gene of interest is cloned downstream of Cerulean, our reporter protein. A second fluorescent protein, mCherry, is incorporated into the same plasmid, to control for transfection efficiency and any non-specific effects of the miR on the transfected cells. The pFmiR plasmid and the miR mimics are cotransfected into Hek-293T cells in a 384-well plate assay. The ratio of Cerulean/mCherry fluorescence in miR transfected cells is normalized to the data from a non-targeting control (NTC) and reflects the extent of miR-target regulation (**Figure 1A**). Our miRFluR assay enables rapid analysis of miR libraries without the need for additional manipulation and reagents post-transfection.

**FIGURE 1.**
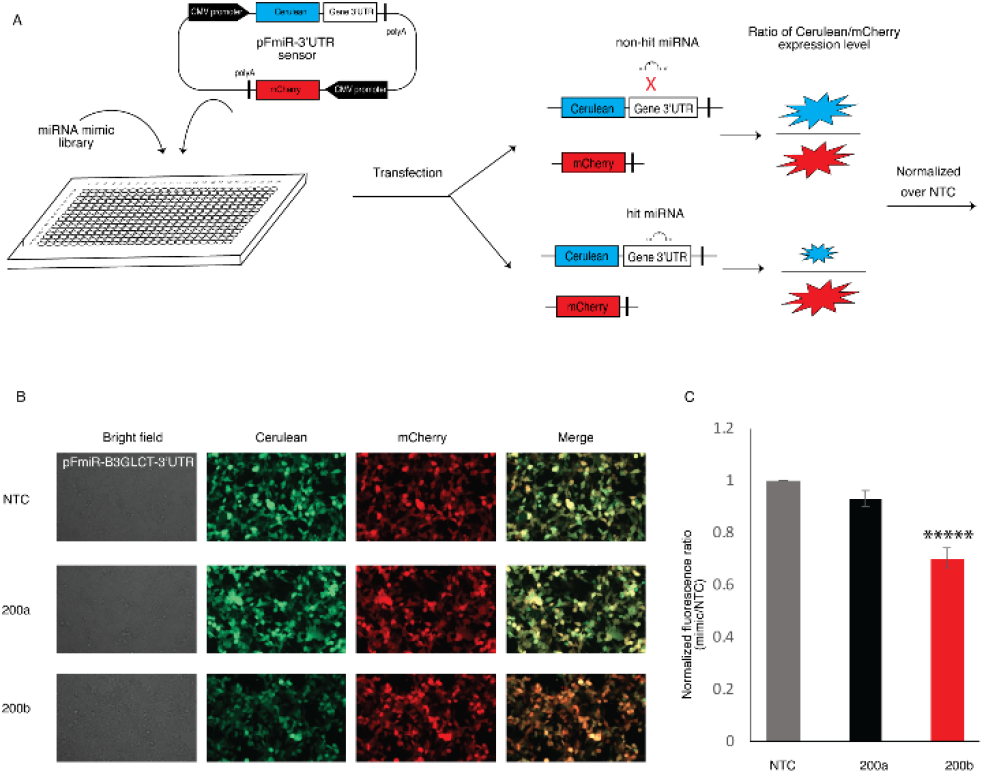
Reproducibility of the miRFluR high-throughput analysis system. (A) Schematic illustration of the miRFluR high-throughput assay. (B) Fluorescence microscopy images of HEK-293T cells co-transfected with pFmiR-B3GLCT and either NTC, miR-200a-3p or miR-200b-3p, 48 h post-transfection. (C) Graphical analysis of data for miR-200a-3p and miR-200b-3p, normalized to NTC, from 27 independent 384-well plates. Unpaired Student’s t test; *****p << 0.0001 (see SI for details).

For our first target, we chose to study the miR regulation of the non-canonical glycosyltranferase, beta-1,3-glucosyltransferase (B3GLCT). This enzyme catalyzes the addition of glucose to *O*-linked fucosyl glycans on thrombospondin type-1 repeats (TSRs) ^13^. Mutations in this gene are known to cause the congenital disorder Peters’ plus syndrome ^18, 19^. In previous work, we identified miR-200b-3p as a regulator of this enzyme and demonstrated a role for B3GLCT in EMT ^10^. Bioinformatic analysis of miRNA predictions identified this glycogene as a highly regulated target ^10^. Thus, we anticipated that a large number of miRs would downregulate this enzyme.

We inserted the 3’UTR of B3GLCT after the stop codon of Cerulean in our pFmiR plasmid to give the pFmiR-B3GLCT-3’UTR sensor (**Figure S2**). To establish that our sensor worked as expected, we compared the Cerulean/mCherry fluorescence ratios upon co-transfection of our sensor with either NTC, the positive control miR-200b-3p or the known negative control miR-200a-3p (**Figure 1B and S3**) ^10^. We observed a clear downregulation of Cerulean, but not mCherry, by miR-200b-3p but not NTC1 or miR-200a-3p. We next analyzed our sensor using the Mission miRNA mimic library v.21 (Sigma), which contained all human miRs included in miRbase version 21. This library has 2,754 miR mimics. We aliquoted these mimics into 384-well plates in triplicate, for a total of 32 plates. Each plate contained NTC, miR-200b-3p and miR-200a-3p as controls. miRs were co-transfected with pFmiR plasmid into Hek-293T cells using lipofectamine 2000. After 48 h, plates were read by a fluorescence plate reader. For each plate, the average ratiometric data for each miR was normalized to the average ratiometric data for the NTC in that plate. Higher error measurements were observed in 5 plates, and these were omitted from further analysis. Comparison of the miR-200a-3p and miR-200b-3p data for the remaining 27 plates showed high reproducibility in the data, with significant repression of B3GLCT observed for miR-200b-3p as compared to miR-200a-3p, in line with our previous work (**Figure 1C**).

### Identification of miR hits for B3GLCT

We next analyzed the remaining miR data for B3GLCT. We first removed any miRs that had high errors in the measurement (median error +2 S.D. across all plates), leaving us with data for 2,071 miRs. We Z-scored the remaining NTC normalized ratiometric data from the sensor. In line with previous work by Wolter et al using luciferase assays ^6^, we set the threshold for hits at 20% change (either up or down) and a Z-score of +/-1.960, which corresponds to the 95% confidence interval. Using these thresholds, we identified 27 miRs that downregulated expression, all of which met the 20% threshold. To our surprise, we also identified 11 miRs that were potential upregulators (**Figure 2A & B**). Although a few upregulatory miRs have been described in the literature ^20, 21^, most are thought to activate expression in senescent cells ^22, 23^. To validate our findings, we first rescreened a small set of 12 miRs (**Figure S4 and S5**). With one exception, all miRs recapitulated the findings observed in the library screen. We next performed Western blot analysis for the protein levels of B3GLCT in HEK-293T transfected with the subset of downregulatory (miR-200b-3p, miR-504, miR-4504, miR-4649-3p, miR-4725-5p) and upregulatory (miR-891b and miR-4470-5p) miRs that passed our secondary screen. We used miR-200a-3p and NTC as negative controls (**Figure 2C-D and S5, Table S1**). In general, the B3GLCT protein levels followed the expected results from our sensor assay, with one exception. The downregulatory miR-4649-3p did not show significant inhibition. We tested whether the mRNA levels of B3GLCT changed with miR transfection (**Figure 2E**). Although generally mRNA levels are thought to correspond to protein expression, the correlation is not absolute and miRs have been shown to impact mRNA in ways not reflected in the protein levels ^6^. For all inhibitory miRs, including miR-4649-3p, we observed a clear loss of mRNA expression for B3GLCT. Conversely, the upregulatory miR, miR-891b, elevated mRNA expression in line with its impact on the protein. Interestingly, miR-4470-5p, which upregulated both protein and sensor expression, clearly repressed mRNA levels for B3GLCT. This argues for multiple pathways to protein regulation through differential mRNA regulation by miRNA.

**FIGURE 2.**
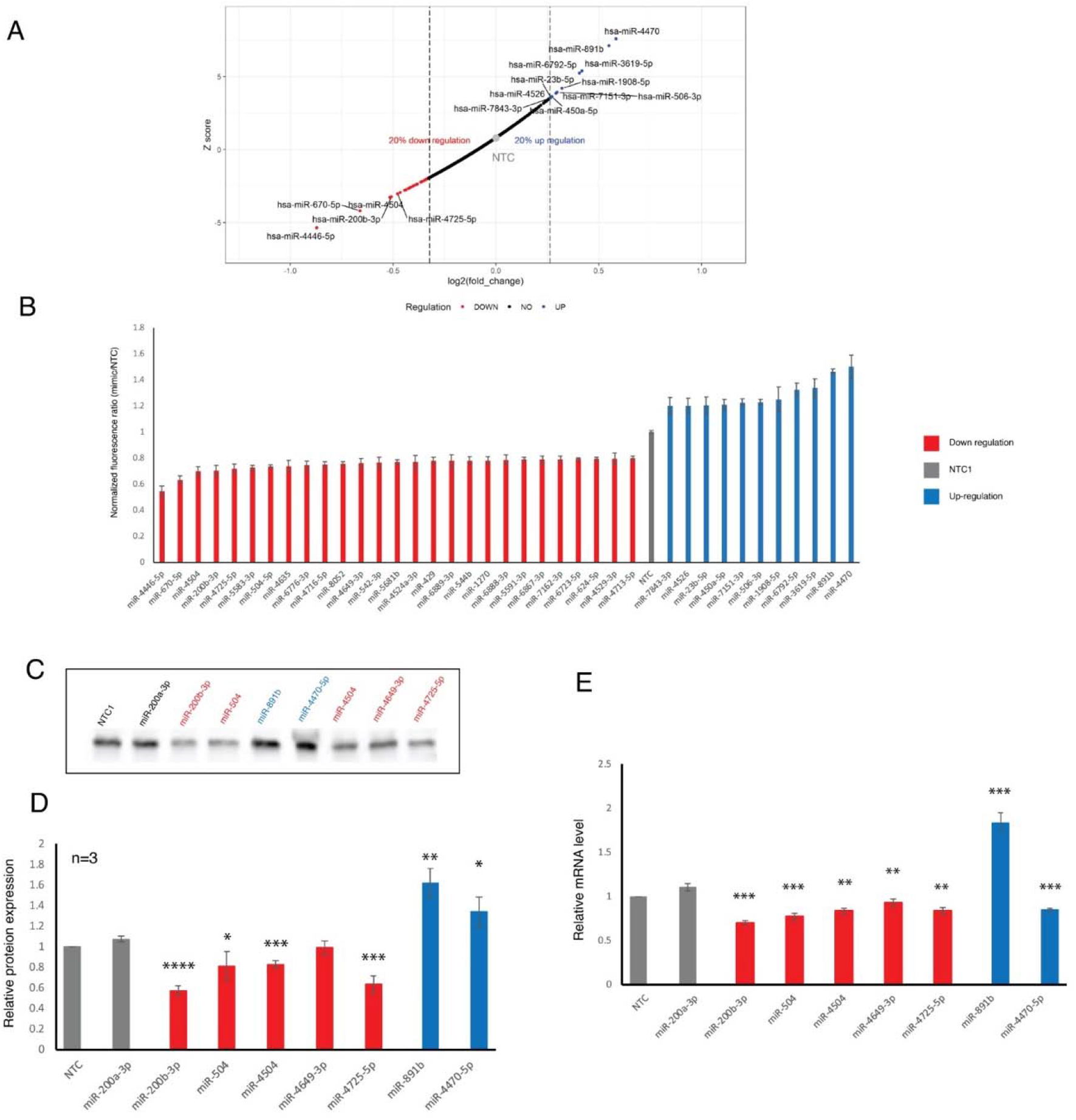
Identification and validation of hits for B3GLCT. (A) Plot of Z-score versus log_2_(fold change) for 2074 miRs against the 3’-UTR of B3GLCT. miRs within the 95 % confidence interval and with a minimum impact of +/− 20 % are labeled (red:downregulatory, blue: upregulatory). NTC is shown in grey for reference. (B) Bar graph of ratiometric data for miRs indicated in A. Error bars represent propagated error. (C) Western blot analysis of B3GLCT in HEK273T transfected with 50 nM miR mimics or NTC, 48 hours posttransfection. (D) Quantification of Western blot analysis for three independent experiments as in C. B3GLCT expression was normalized to total protein and set over normalized NTC for each blot. Statistical analysis was done against miR-200a-3p as a negative control. (E) RT-qPCR analysis for relative B3GLCT mRNA expression levels. All samples are normalized to GAPDH within the sample and then to NTC for that run. Results shown are from three independent experiments. Statistical analysis was done against miR-200a-3p as a negative control. Student’s t-test; \**p*<0.05, ** *p* <0.01, *** *p* <0.001 and **** *p* <0.0001.

### Comparison of prediction to experimental data for miR regulation of B3GLCT

Identification of miRs that target a specific protein is heavily based on prediction from algorithms. We tested how accurately two of the most popular miR prediction programs, Targetscan 7.2 ^3^ and miRwalk 3.0 ^4, 24, 25^, predicted B3GLCT regulators. For both algorithms, we only examined miR predictions for miR within our final dataset. Targetscan 7.2 predicted 480 unique miR interactions with the 3’-UTR of B3GLCT (**Figure 3A**). Of those, only 17 (3.5%) were identified as hits within our screen. All 17 were repressors. Of the repressors, 17/27 (~2/3) were identified by Targetscan.

**FIGURE 3.**
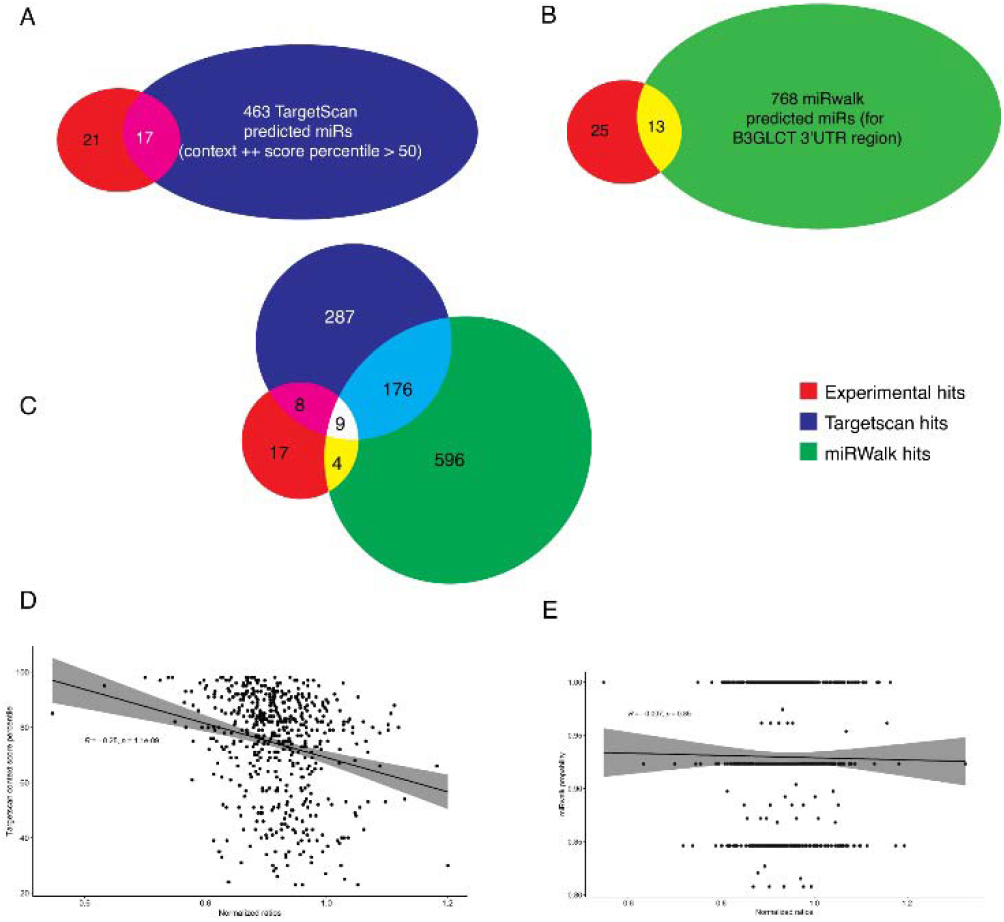
Comparison of experimental results to prediction datasets from TargetScan (A) or miRWalk (B). (C) Overlap of experimental results and predictions from both Targetscan and miRWalk. (D) Correlation between Targetscan context++ score percentile and our experimental results. A significant but small negative correlation was observed (R = −0.25, *p*~10^-9^) with data for which Targetscan context scores++ exist. (E) Correlation between miRWalk score and our experimental results. No correlation was observed.

Overall, there was a weak but significant correlation between the Targetscan score, where available, and the level of repression observed (**Figure 3D,** R= 0.25, *p* = 1 x 10^-9^). It should be noted, however, that although Targetscan 7.2 analyzes the miRs from miRbase v 21, only 559 of the 2,071 miRs from our analysis have a context score in Targetscan. Scores were not available for the 10 non-predicted downregulatory miRs or the 10 upregulatory miRs. Among the unpredicted downregulators were miR-504-5p, miR-4649-3p and miR-4725-5p, all of which showed clear repression of B3GLCT in our assays (**Figure 2 C-E**). Thus, the actual correlation is likely far lower. For miRwalk 3.0, 781 unique miR interactions were predicted. Of these, only 13 were observed (1.7%, **Figure 3B**). In this case, 1 of the upregulators (miR-6792-5p) was among the predicted hits. No correlation was observed between the score in miRwalk 3.0 and miR regulation of the sensor (**Figure 3E**). Only 9 of the hits were predicted by both algorithms, which only predicted 185 miRs in common between the two (**Figure 3C**). In previous work, a higher concordance between prediction and testing (~16^7^-63 %^6^) was observed. However, in that work, multiple 3’-UTRs were tested against a limited set of miRs. These datasets were skewed towards highly abundant miRs and cancer-related genes. Our analysis is the first to test a broad swath of the human miRome against a single gene, unrelated to cancer. Our approach shows that current prediction algorithms are significantly less accurate than previously thought, with a high bias towards false positives.

### miRs Downregulating B3GLCT Predict Peters’ Plus Syndrome

In previous work we posited that miR regulation of a gene would predict the biological functions of that gene, a hypothesis we termed the miRNA Proxy Hypothesis ^8, 10, 11^. Downregulation of B3GLCT activity has a known set of biological outcomes due to the existence of mutations in the gene in the human population, which cause a loss of activity. These result in the genetic disorder Peters’ Plus Syndrome, which is characterized by a variety of symptoms including ocular abnormalities, short stature, cleft palate, small ears, facial abnormalities and brain abnormalities, including intellectual disability ^18, 19^. To test our hypothesis, we tested whether miRs that downregulate B3GLCT are predictive of the Peters’ Plus Syndrome phenotypes. We analyzed the gene target network, and enrichment in associated disease phenotypes, of our 27 validated downregulatory miRs using miRNet ^26, 27^. Only validated miR-target interactions from miRTarbase were used in our evaluation. None of the 27 miRs fed into the system were known in miRTarbase to target B3GLCT. The gene target network for our downregulatory miRs was functionally enriched in genes associated with Peters’ Plus Syndrome features (**Figure 4, Table S2**). The only non-Peters’ plus phenotype observed in the predicted set were a subset of cancer phenotypes. Whether this is due to a real role for B3GLCT in these specific cancers, or is an outcome of the bias of the current datasets towards cancer genes is unknown. Overall, our analysis supports the miRNA proxy hypothesis, predicting a role for B3GLCT in the disease outcomes related to Peters’ Plus Syndrome through the miRs that downregulate this enzyme.

**FIGURE 4.**
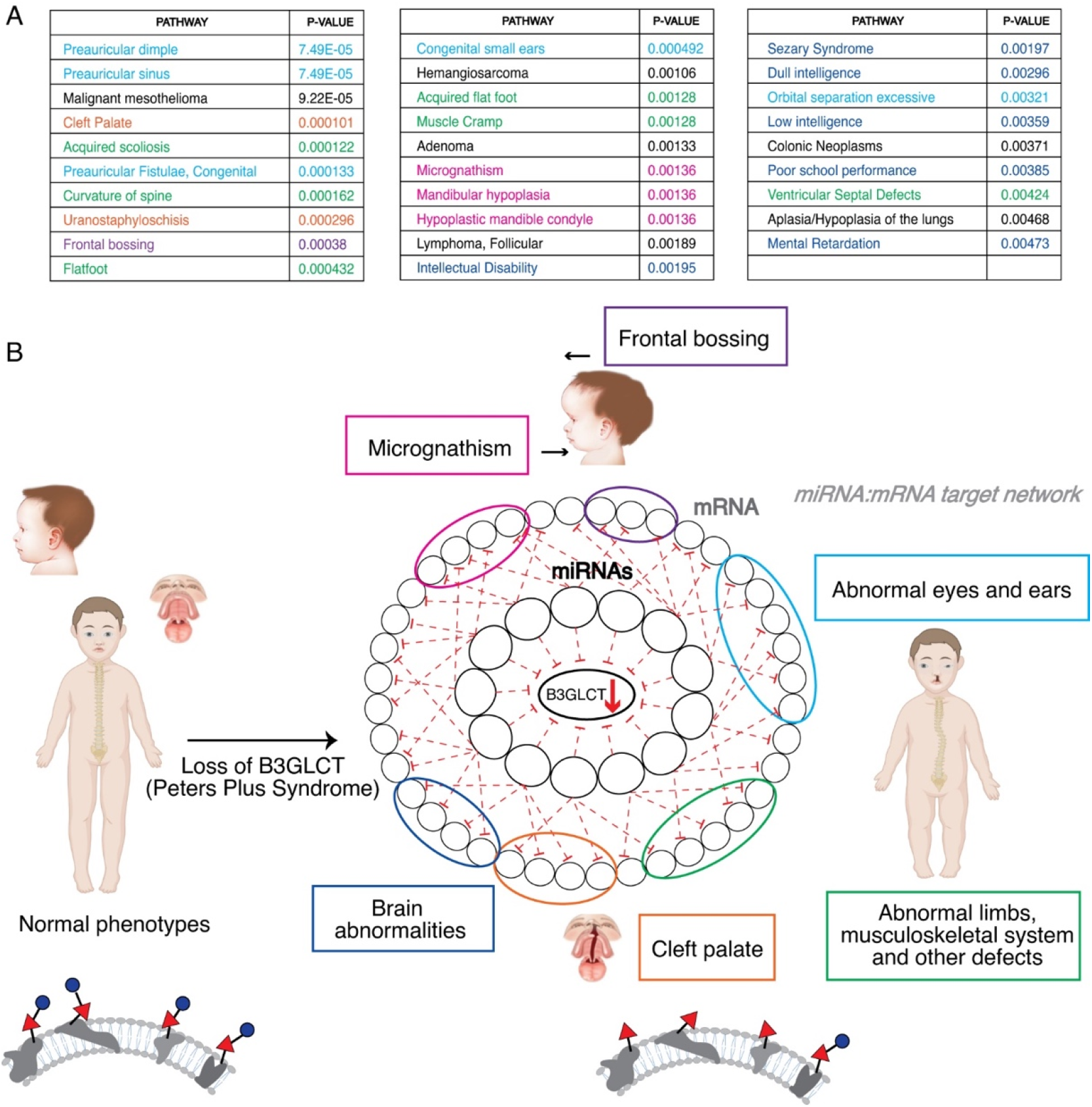
Phenotypic network analysis of miRs downregulating B3GLCT. (A) Table of enriched disease phenotypes resulting from miRNet analysis of B3GLCT downregulatory miRs. Table is color coded to phenotypes seen in Peter’s Plus Syndrome as in (B). (B) Schematic of B3GLCT downregulating miR-mRNA target network as it applies identification of disease phenotypes observed in Peters’ Plus Syndrome. miRs that downregulate B3GLCT target the mRNA of genes enriched in the disease networks shown in (A).

## CONCLUSIONS

Our current understanding of miR regulation of protein expression has been hampered by limited data on miR:mRNA target interactions. Herein, we created a high-throughput experimental platform, miRFluR, to rapidly analyze miR interactions with the 3’-UTR of a gene of interest. We used this dual fluorescence platform to perform the first comprehensive analysis of miR regulation of a gene, B3GLCT, through its 3’-UTR. Our analysis found both downregulatory and upregulatory miRs for B3GLCT, which we validated at the protein and mRNA levels. We anticipated that this gene would be highly regulated, based on the predictions from multiple algorithms ^3, 10, 28^. However, we found a wide discrepancy between prediction and our assay, with < 4% of predicted miRs targeting this enzyme (>96 % false positive rate). Although it is widely held that miRs target hundreds to thousands of genes, our results would argue that prediction algorithms vastly overstate miR regulation. Functional enrichment analysis of miRs downregulating B3GLCT identified disease phenotypes included in Peters’ Plus Syndrome, the known disorder caused by mutation of this gene, in line with our miRNA Proxy hypothesis. One limitation of this analysis is that the dataset underpinning miRNet and other such network analysis algorithms has a lack of validated interactions^5, 29^. As our information on true miR:target interactions grow, our ability to harness this data to understand the biological functions of the glycome and other genes will improve.

## EXPERIMENTAL METHODS

### Cloning of pFmiR-B3GLCT-3’UTR

B3GLCT 3’UTR was cloned from cDNA using primers:

B3GLCTc_fwd: CTAGCATCAGGGTGACCTG

B3GLCT_rev: GATCCTTTTCATTACATAATAAAG

and standard PCR conditions. The DNA fragment was cloned using the NheI and BamHI sites downstream of Cerulean in our pFmiR-empty backbone using standard ligation protocols. Plasmid maps and sequences for pFmiR and pFmiR-B3GLCT-3’-UTR can be found in **Figure S1** and **Figure S2**, respectively.

### FluoRmiR High-throughput Assay

The Human miRNA Mimic library version 21 (MISSION, Sigma) was resuspended in nuclease-free water and aliquoted into black 384-well, clear optical bottom tissue-culture treated plates (Nunc). Each plate contained 3 replicates of every miRNA (1.8 pmol/well). Including controls (NTC, miR-200a-3p, miR-200b-3p).

To each well in the plate was added 25 ng of pMIR-B3GLCT plasmid in 5 μl Opti-MEM (Gibco) and 0.11 μl lipofectamine 2000 (Invitrogen) in 5 μl Opti-MEM (Gibco). The solution was allowed to incubate at room temperature for 25 min. Then, HEK293T cells (25 μl per well, 400 cells/ μl in non-phenol red Dulbecco’s Modified Eagle Medium (DMEM) with FBS 10%) were added to the plate. Plates were then incubated at 37°C, 5% CO_2_. After 48 hours, the fluorescence signals of Cerulean (excitation: 433 nm; emission: 475 nm) and mCherry (excitation: 587 nm; emission: 610 nm) were measured using the bottom read option in a FlexStation 3 Multi-mode microplate reader (Molecular Devices).

### Data Processing

We calculated the ratio Cerulean fluorescence (Cer) over mCherry fluorescence (Cer/mCh) for each well in each plate. For each miR, triplicate values were averaged and the standard deviation (S.D.) obtained. We calculated a % error for each miR as 100 x S.D./mean.. As a quality control measure, we removed any plates or miRs that had high errors in the measurement (median error +2 S.D. across all plates). This left us with data for 2,071 miRs. The Cer/mCh ratio for each miR was then normalized to the Cer/mCh ratio for the NTC within that plate and error was propagated. Data from all plates was then combined and Z-scores were calculated. A Z-score of +/-1.960, corresponding to a 2-tailed p-value of 0.05, was used as a threshold for significance. In addition, we set a second threshold of +/− 20% impact by the miR, in line with previous work ^6, 7^.

### Western Blot

HEK293T cells were seeded in six-well plates (80,000 cells per well), cultured for 24 h, and transfected with miRNA mimics (50 nM, Sigma MISSION) using Lipofectamine 2000 (Life Technologies). Cells were washed and harvested 48 hours post-transfection.

Cells were then lysed in cold RIPA buffer supplemented with protease inhibitors and 50 μg of protein were run on SDS-PAGE. Standard Western Blot analysis using α-B3GLCT (IHC-plus antihuman B3GALTL antibody, 1:500) and α-rabbit-HRP (2°, 1:5,000, Abcam)] was performed ^9^. Blots were developed using Clarity and Clarity Max Western ECL substrate (Bio-Rad).

### RT-PCR

Total RNA was isolated using RNeasy kit (Qiagen) according to the manufacturer’s instructions. RNA concentrations were measured using NanoDrop, and isolated RNA was reverse-transcribed (Applied Biosystems Power SYBR Green PCR). Realtime quantitative PCR (qPCR) was performed using the SYBR Green method and cycle threshold values (Ct) were obtained using an Applied Biosystem (ABI) 7500 Real-Time PCR machine and normalized to GAPDH.

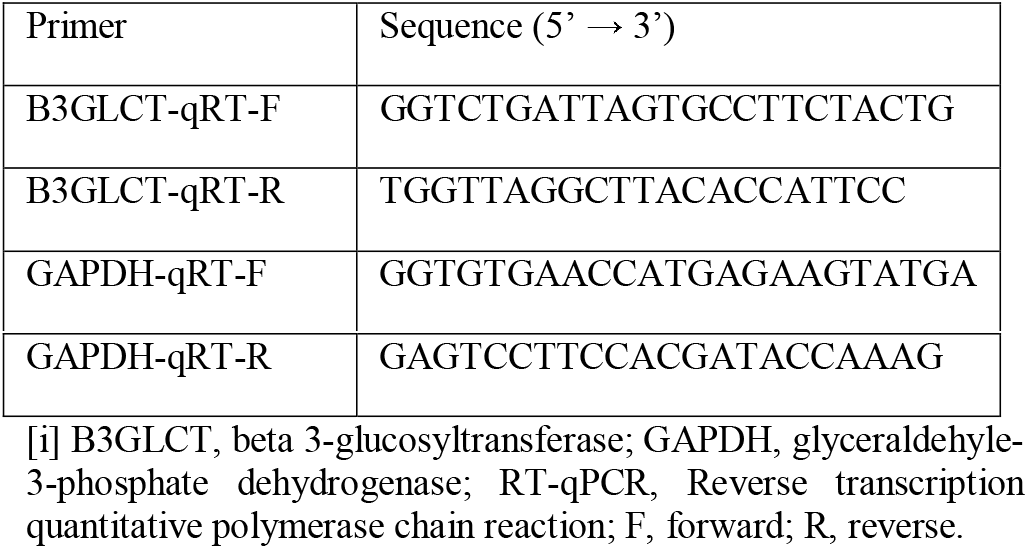

### Network Analysis Using miRNet

Downregulatory miRs (27 miRs, Figure 2B) were input into miRNet (www.mirnet.ca) ^26,27^ using the following parameters: Organism: human miRs, ID type: miRbase ID, Targets: Genes(miRTarbase v 8.0). The Diseases Phenotype Enrichment function was used for Figure 4.

## ASSOCIATED CONTENT

### Supporting Information

Supporting information and figures (PDF)

Supplementary tables (XLSX)

## AUTHOR INFORMATION

### Author Contributions

**Chu T. Thu** - *Department of Chemistry, University of Alberta, Edmonton, T, Canada;* https://orcid.org/0000-0001-5172-3952;

**Jonathan Y. Chung** - *Department of Chemistry, New York University, New York 10003, United States;*

**Deepika Dhawan** - *Department of Chemistry, New York University, New York 10003, United States;*

**Christopher Vaiana** - *Department of Chemistry, New York University, New York 10003, United States;*

### Notes

The authors declare no competing financial interests.

## ACKNOWLEDGMENT

This work was supported by the National Institutes of Health Common Fund (Grant U01CA221229), the Department of Defense (DARPA-BAA-16-51) and the Canada Excellence Research Chairs Program.

---

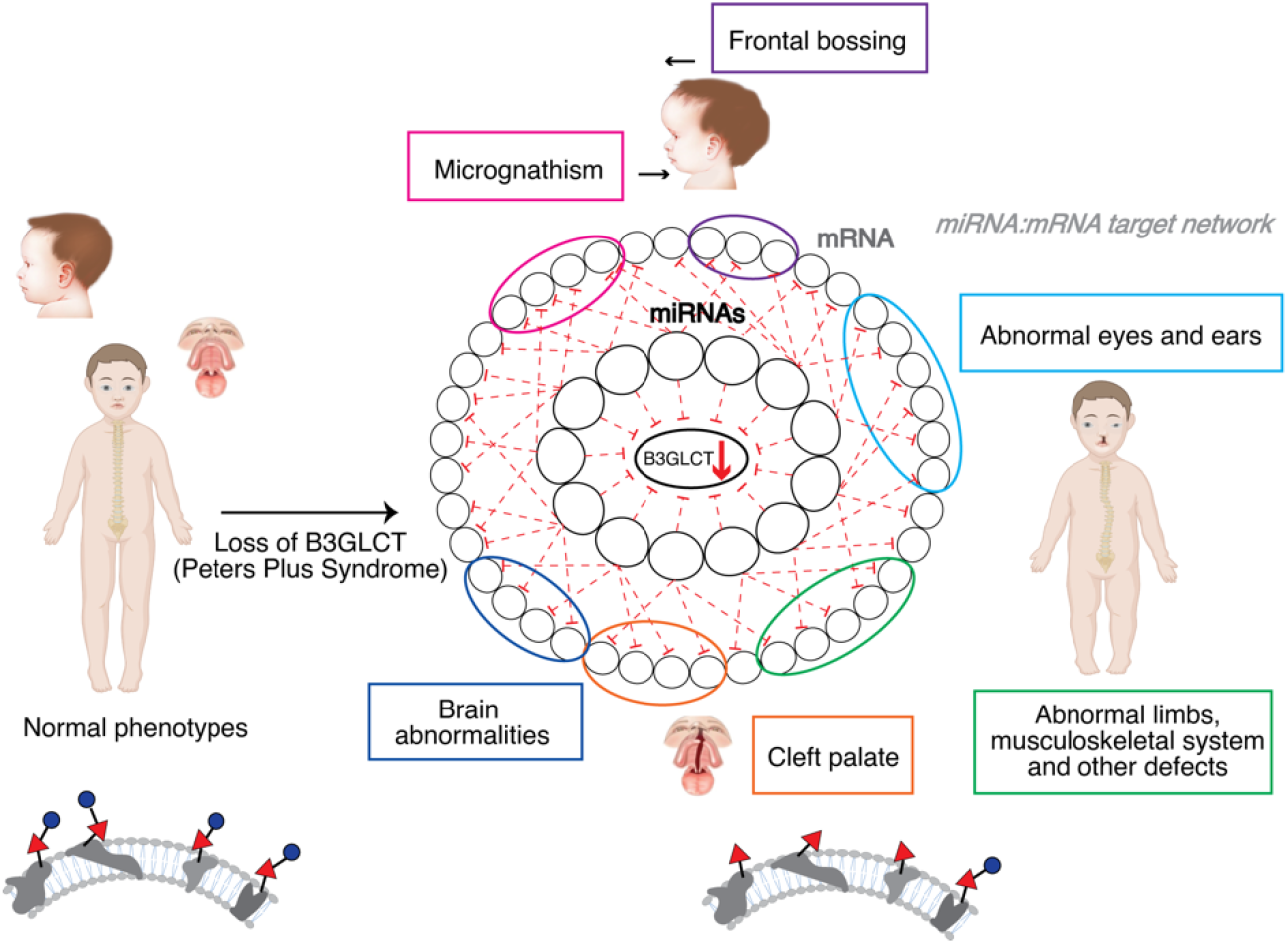

## REFERENCES

(1) Schmiedel, J. M.; Klemm, S. L.; Zheng, Y.; Sahay, A.; Bluthgen, N.; Marks, D. S.; van Oudenaarden, A., Gene expression. MicroRNA controlof protein expression noise. Science 2015, 348 (6230), 128–32.

(2) Liu, B.; Li, J.; Cairns, M. J., Identifying miRNAs, targets and functions. Brief Bioinform 2014, 15 (1), 1–19.

(3) Agarwal, V.; Bell, G. W.; Nam, J. W.; Bartel, D. P., Predicting effective microRNA target sites in mammalian mRNAs. Elife 2015, 4.

(4) Dweep, H.; Sticht, C.; Pandey, P.; Gretz, N., miRWalk--database: prediction of possible miRNA binding sites by “walking”the genes of three genomes. J Biomed Inform 2011, 44 (5), 839–47.

(5) Hsu, S. D.; Lin, F. M.; Wu, W. Y.; Liang, C.; Huang, W. C.; Chan, W. L.; Tsai, W. T.; Chen, G. Z.; Lee, C. J.; Chiu, C. M.; Chien, C. H.; Wu, M. C.; Huang, C. Y.; Tsou, A. P.; Huang, H. D., miRTarBase: a database curates experimentally validated microRNA-target interactions. Nucleic Acids Res 2011, 39 (Database issue), D163–9.

(6) Wolter, J. M.; Kotagama, K.; Pierre-Bez, A. C.; Firago, M.; Mangone, M., 3’LIFE: a functional assay to detect miRNA targets in high-throughput. Nucleic Acids Res 2014, 42 (17), e132.

(7) Zhou, P.; Xu, W.; Peng, X.; Luo, Z.; Xing, Q.; Chen, X.; Hou, C.; Liang, W.; Zhou, J.; Wu, X.; Songyang, Z.; Jiang, S., Large-scale screens of miRNA-mRNA interactions unveiled that the 3’UTR of a gene is targeted by multiple miRNAs. PLoS One 2013, 8 (7), e68204.

(8) Thu, C. T.; Mahal, L. K., Sweet Control: MicroRNA Regulation of the Glycome. Biochemistry 2020, 59 (34), 3098–3110.

(9) Agrawal, P.; Kurcon, T.; Pilobello, K. T.; Rakus, J. F.; Koppolu, S.; Liu, Z.; Batista, B. S.; Eng, W. S.; Hsu, K. L.; Liang, Y.; Mahal, L. K., Mapping posttranscriptional regulation of the human glycome uncovers microRNA defining the glycocode. Proc Natl Acad Sci USA 2014, 111 (11), 4338–43.

(10) Kurcon, T.; Liu, Z.; Paradkar, A. V.; Vaiana, C. A.; Koppolu, S.; Agrawal, P.; Mahal, L. K., miRNA proxy approach reveals hidden functions of glycosylation. Proc Natl Acad Sci U SA 2015, 112 (23), 7327–32.

(11) Vaiana, C. A.; Kurcon, T.; Mahal, L. K., MicroRNA-424 Predicts a Role for beta-1,4 Branched Glycosylation in Cell Cycle Progression. J Biol Chem 2016, 291 (3), 1529–37.

(12) Varki, A., Biological roles of glycans. Glycobiology 2017, 27 (1), 3–49.

(13) Holdener, B. C.; Percival, C. J.; Grady, R. C.; Cameron, D. C.; Berardinelli, S. J.; Zhang, A.; Neupane, S.; Takeuchi, M.; Jimenez-Vega, J. C.; Uddin, S. M. Z.; Komatsu, D. E.; Honkanen, R.; Dubail, J.; Apte, S. S.; Sato, T.; Narimatsu, H.; McClain, S. A.; Haltiwanger, R. S., ADAMTS9 and ADAMTS20 are differentially affected by loss of B3GLCT in mouse model of Peters plus syndrome. Hum Mol Genet 2019, 28 (24), 4053–4066.

(14) Chi, S. W.; Zang, J. B.; Mele, A.; Darnell, R. B., Argonaute HITS-CLIP decodes microRNA-mRNA interaction maps. Nature 2009, 460(7254), 479–86.

(15) Bracken, C. P.; Li, X.; Wright, J. A.; Lawrence, D. M.; Pillman, K. A.; Salmanidis, M.; Anderson, M. A.; Dredge, B. K.; Gregory, P. A.; Tsykin, A.; Neilsen, C.; Thomson, D. W.; Bert, A. G.; Leerberg, J. M.; Yap, A. S.; Jensen, K. B.; Khew-Goodall, Y.; Goodall, G. J., Genomewide identification of miR-200 targets reveals a regulatory network controlling cell invasion. EMBO J 2014, 33 (18), 2040–56.

(16) Mukherji, S.; Ebert, M. S.; Zheng, G. X.; Tsang, J. S.; Sharp, P. A.; van Oudenaarden, A., MicroRNAs can generate thresholds in target gene expression. Nat Genet 2011, 43 (9), 854–9.

(17) Lemus-Diaz, N.; Boker, K. O.; Rodriguez-Polo, I.; Mitter, M.; Preis, J.; Arlt, M.; Gruber, J., Dissecting miRNA gene repression on single cell level with an advanced fluorescent reporter system. Sci Rep 2017, 7, 45197.

(18) Maillette de Buy Wenniger-Prick, L. J.; Hennekam, R. C., The Peters’ plus syndrome: a review. Ann Genet 2002, 45 (2), 97–103.

(19) Hennekam, R. C.; Van Schooneveld, M. J.; Ardinger, H. H.; Van Den Boogaard, M. J.; Friedburg, D.; Rudnik-Schoneborn, S.; Seguin, J. H.; Weatherstone, K. B.; Wittebol-Post, D.; Meinecke, P., The Peters’-Plus syndrome: description of 16 patients and review of the literature. Clin Dysmorphol 1993, 2 (4), 283–300.

(20) Liu, J.; Li, Y.; Chen, X.; Xu, X.; Zhao, H.; Wang, S.; Hao, J.; He, B.; Liu, S.; Wang, J., Upregulation of miR-205 induces CHN1 expression, which is associated with the aggressive behaviour of cervical cancer cells and correlated with lymph node metastasis. BMC Cancer 2020, 20 (1), 1029.

(21) Wang, L.; Zhang, S.; Zhang, W.; Cheng, G.; Khan, R.; Junjvlieke, Z.; Li, S.; Zan, L., miR-424 Promotes Bovine Adipogenesis Through an Unconventional Post-Transcriptional Regulation of STK11. Front Genet 2020, 11, 145.

(22) Vasudevan, S.; Tong, Y.; Steitz, J. A., Switching from repression to activation: microRNAs can up-regulate translation. Science 2007, 318 (5858), 1931–4.

(23) Valinezhad Orang, A.; Safaralizadeh, R.; Kazemzadeh-Bavili, M., Mechanisms of miRNA-Mediated Gene Regulation from Common Downregulation to mRNA-Specific Upregulation. Int J Genomics 2014, 2014, 970607.

(24) Dweep, H.; Gretz, N.; Sticht, C., miRWalk database for miRNA-target interactions. Methods Mol Biol 2014, 1182, 289–;305.

(25) Sticht, C.; De La Torre, C.; Parveen, A.; Gretz, N., miRWalk: An online resource for prediction of microRNA binding sites. PLoS One 2018, 13 (10), e0206239.

(26) Fan, Y.; Siklenka, K.; Arora, S. K.; Ribeiro, P.; Kimmins, S.; Xia, J., miRNet - dissecting miRNA-target interactions and functional associations through network-based visual analysis. Nucleic Acids Res 2016, 44 (W1), W135–41.

(27) Chang, L.; Zhou, G.; Soufan, O.; Xia, J., miRNet 2.0: networkbased visual analytics for miRNA functional analysis and systems biology. Nucleic Acids Res 2020, 48 (W1), W244–W251.

(28) Dweep, H.; Gretz, N., miRWalk2.0: a comprehensive atlas of microRNA-target interactions. Nat Methods 2015, 12 (8), 697.

(29) Huang, H. Y.; Lin, Y. C.; Li, J.; Huang, K. Y.; Shrestha, S.; Hong, H. C.; Tang, Y.; Chen, Y. G.; Jin, C. N.; Yu, Y.; Xu, J. T.; Li, Y. M.; Cai, X. X.; Zhou, Z. Y.; Chen, X. H.; Pei, Y. Y.; Hu, L.; Su, J. J.; Cui, S. D.; Wang, F.; Xie, Y. Y.; Ding, S. Y.; Luo, M. F.; Chou, C. H.; Chang, N. W.; Chen, K. W.; Cheng, Y. H.; Wan, X. H.; Hsu, W. L.; Lee, T. Y.; Wei, F. X.; Huang, H. D., miRTarBase 2020: updates to the experimentally validated microRNA-target interaction database. Nucleic Acids Res 2020, 48 (D1), D148–D154.

