## Supplementary Information for "High-throughput analysis of B3GLCT regulation predicts phenotype of Peters’ Plus Syndrome in line with the miRNA Proxy Hypothesis"

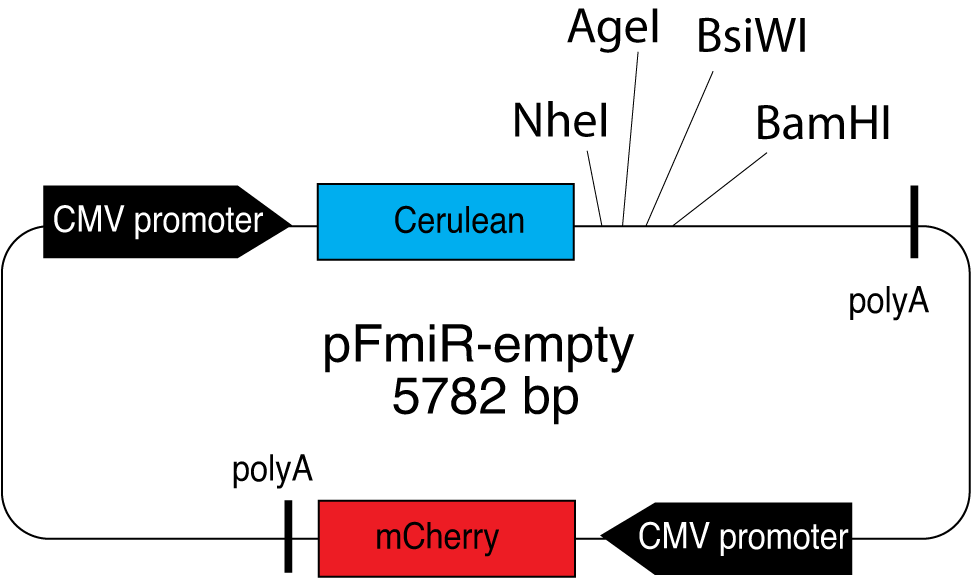

List of features:

mCherry (2539..3249)

CMV promoter 1 (236..852)

CMV promoter 2 (1916..2532)

Cerulean (918..1637)

polyA_1 (1688..1914)

polyA_2 (3272..3498)

Sequence:

GACGGATCGGGAGATCTCCCGATCCCCTATGGTCGACTCTCAGTACAATCTGCTCTGATGCCGCATAGTTAAGCCAGTATCTGCTCCCTGCTTGTGTGTTGGAGGTCGCTGAGTAGTGCGCGAGCAAAATTTAAGCTACAACAAGGCAAGGCTTGACCGACAATTGCATGAAGAATCTGCTTAGGGTTAGGCGTTTTGCGCTGCTTCGCGATGTACGGGCCAGATATACGCGTTGACATTGATTATTGACTAGTTATTAATAGTAATCAATTACGGGGTCATTAGTTCATAGCCCATATATGGAGTTCCGCGTTACATAACTTACGGTAAATGGCCCGCCTGGCTGACCGCCCAACGACCCCCGCCCATTGACGTCAATAATGACGTATGTTCCCATAGTAACGCCAATAGGGACTTTCCATTGACGTCAATGGGTGGACTATTTACGGTAAACTGCCCACTTGGCAGTACATCAAGTGTATCATATGCCAAGTACGCCCCCTATTGACGTCAATGACGGTAAATGGCCCGCCTGGCATTATGCCCAGTACATGACCTTATGGGACTTTCCTACTTGGCAGTACATCTACGTATTAGTCATCGCTATTACCATGGTGATGCGGTTTTGGCAGTACATCAATGGGCGTGGATAGCGGTTTGACTCACGGGGATTTCCAAGTCTCCACCCCATTGACGTCAATGGGAGTTTGTTTTGGCACCAAAATCAACGGGACTTTCCAAAATGTCGTAACAACTCCGCCCCATTGACGCAAATGGGCGGTAGGCGTGTACGGTGGGAGGTCTATATAAGCAGAGCTCTCTGGCTAACTAGAGAACCCACTGCTTACTGGCTTATCGAAATTAATACGACTCACTATAGGGAGACCCAAGCTTGGTACCGAGCTCGGATCGATATCATGGTGAGCAAGGGCGAGGAGCTGTTCACCGGGGTGGTGCCCATCCTGGTCGAGCTGGACGGCGACGTAAACGGCCACAAGTTCAGCGTGTCCGGCGAGGGCGAGGGCGATGCCACCTACGGCAAGCTGACCCTGAAGTTCATCTGCACCACCGGCAAGCTGCCCGTGCCCTGGCCCACCCTCGTGACCACCCTGACCTGGGGCGTGCAGTGCTTCGCCCGCTACCCCGACCACATGAAGCAGCACGACTTCTTCAAGTCCGCCATGCCCGAAGGCTACGTCCAGGAGCGCACCATCTTCTTCAAGGACGACGGCAACTACAAGACCCGCGCCGAGGTGAAGTTCGAGGGCGACACCCTGGTGAACCGCATCGAGCTGAAGGGCATCGACTTCAAGGAGGACGGCAACATCCTGGGGCACAAGCTGGAGTACAACGCCATCAGCGACAACGTCTATATCACCGCCGACAAGCAGAAGAACGGCATCAAGGCCAACTTCAAGATCCGCCACAACATCGAGGACGGCAGCGTGCAGCTCGCCGACCACTACCAGCAGAACACCCCCATCGGCGACGGCCCCGTGCTGCTGCCCGACAACCACTACCTGAGCACCCAGTCCGCCCTGAGCAAAGACCCCAACGAGAAGCGCGATCACATGGTCCTGCTGGAGTTCGTGACCGCCGCCGGGATCACTCTCGGCATGGACGAGCTGTACAAGTAATAAGCTAGCACACCGGTCGTCTAGCAATGCGATCCACGTACGTAGGATCCCGACTGTGCCTTCTAGTTGCCAGCCATCTGTTGTTTGCCCCTCCCCCGTGCCTTCCTTGACCCTGGAAGGTGCCACTCCCACTGTCCTTTCCTAATAAAATGAGGAAATTGCATCGCATTGTCTGAGTAGGTGTCATTCTATTCTGGGGGGTGGGGTGGGGCAGGACAGCAAGGGGGAGGATTGGGAAGACAATAGCAGGCATGCTGGGGATGCGGTGGGCTCTATGGACATTGATTATTGACTAGTTATTAATAGTAATCAATTACGGGGTCATTAGTTCATAGCCCATATATGGAGTTCCGCGTTACATAACTTACGGTAAATGGCCCGCCTGGCTGACCGCCCAACGACCCCCGCCCATTGACGTCAATAATGACGTATGTTCCCATAGTAACGCCAATAGGGACTTTCCATTGACGTCAATGGGTGGACTATTTACGGTAAACTGCCCACTTGGCAGTACATCAAGTGTATCATATGCCAAGTACGCCCCCTATTGACGTCAATGACGGTAAATGGCCCGCCTGGCATTATGCCCAGTACATGACCTTATGGGACTTTCCTACTTGGCAGTACATCTACGTATTAGTCATCGCTATTACCATGGTGATGCGGTTTTGGCAGTACATCAATGGGCGTGGATAGCGGTTTGACTCACGGGGATTTCCAAGTCTCCACCCCATTGACGTCAATGGGAGTTTGTTTTGGCACCAAAATCAACGGGACTTTCCAAAATGTCGTAACAACTCCGCCCCATTGACGCAAATGGGCGGTAGGCGTGTACGGTGGGAGGTCTATATAAGCAGAGCTCTCTGGCTAACTAGAGAACCCACTGCTTACTGGCCCCGGGATGGTGAGCAAGGGCGAGGAGGATAACATGGCCATCATCAAGGAGTTCATGCGCTTCAAGGTGCACATGGAGGGCTCCGTGAACGGCCACGAGTTCGAGATCGAGGGCGAGGGCGAGGGCCGCCCCTACGAGGGCACCCAGACCGCCAAGCTGAAGGTGACCAAGGGTGGCCCCCTGCCCTTCGCCTGGGACATCCTGTCCCCTCAGTTCATGTACGGCTCCAAGGCCTACGTGAAGCACCCCGCCGACATCCCCGACTACTTGAAGCTGTCCTTCCCCGAGGGCTTCAAGTGGGAGCGCGTGATGAACTTCGAGGACGGCGGCGTGGTGACCGTGACCCAGGACTCCTCCCTGCAGGACGGCGAGTTCATCTACAAGGTGAAGCTGCGCGGCACCAACTTCCCCTCCGACGGCCCCGTAATGCAGAAGAAGACCATGGGCTGGGAGGCCTCCTCCGAGCGGATGTACCCCGAGGACGGCGCCCTGAAGGGCGAGATCAAGCAGAGGCTGAAGCTGAAGGACGGCGGCCACTACGACGCTGAGGTCAAGACCACCTACAAGGCCAAGAAGCCCGTGCAGCTGCCCGGCGCCTACAACGTCAACATCAAGTTGGACATCACCTCCCACAACGAGGACTACACCATCGTGGAACAGTACGAACGCGCCGAGGGCCGCCACTCCACCGGCGGCATGGACGAGCTGTACAAGTAATCTAGAGCTCGCTGATCAGCCTCGACTGTGCCTTCTAGTTGCCAGCCATCTGTTGTTTGCCCCTCCCCCGTGCCTTCCTTGACCCTGGAAGGTGCCACTCCCACTGTCCTTTCCTAATAAAATGAGGAAATTGCATCGCATTGTCTGAGTAGGTGTCATTCTATTCTGGGGGGTGGGGTGGGGCAGGACAGCAAGGGGGAGGATTGGGAAGACAATAGCAGGCATGCTGGGGATGCGGTGGGCTCTATGGCTTCTGAGGCGGAAAGAACCAGCTGGGGCTCGAGTGCATTCTAGTTGTGGTTTGTCCAAACTCATCAATGTATCTTATCATGTCTGTATACCGTCGACCTCTAGCTAGAGCTTGGCGTAATCATGGTCATAGCTGTTTCCTGTGTGAAATTGTTATCCGCTCACAATTCCACACAACATACGAGCCGGAAGCATAAAGTGTAAAGCCTGGGGTGCCTAATGAGTGAGCTAACTCACATTAATTGCGTTGCGCTCACTGCCCGCTTTCCAGTCGGGAAACCTGTCGTGCCAGCTGCATTAATGAATCGGCCAACGCGCGGGGAGAGGCGGTTTGCGTATTGGGCGCTCTTCCGCTTCCTCGCTCACTGACTCGCTGCGCTCGGTCGTTCGGCTGCGGCGAGCGGTATCAGCTCACTCAAAGGCGGTAATACGGTTATCCACAGAATCAGGGGATAACGCAGGAAAGAACATGTGAGCAAAAGGCCAGCAAAAGGCCAGGAACCGTAAAAAGGCCGCGTTGCTGGCGTTTTTCCATAGGCTCCGCCCCCCTGACGAGCATCACAAAAATCGACGCTCAAGTCAGAGGTGGCGAAACCCGACAGGACTATAAAGATACCAGGCGTTTCCCCCTGGAAGCTCCCTCGTGCGCTCTCCTGTTCCGACCCTGCCGCTTACCGGATACCTGTCCGCCTTTCTCCCTTCGGGAAGCGTGGCGCTTTCTCAATGCTCACGCTGTAGGTATCTCAGTTCGGTGTAGGTCGTTCGCTCCAAGCTGGGCTGTGTGCACGAACCCCCCGTTCAGCCCGACCGCTGCGCCTTATCCGGTAACTATCGTCTTGAGTCCAACCCGGTAAGACACGACTTATCGCCACTGGCAGCAGCCACTGGTAACAGGATTAGCAGAGCGAGGTATGTAGGCGGTGCTACAGAGTTCTTGAAGTGGTGGCCTAACTACGGCTACACTAGAAGGACAGTATTTGGTATCTGCGCTCTGCTGAAGCCAGTTACCTTCGGAAAAAGAGTTGGTAGCTCTTGATCCGGCAAACAAACCACCGCTGGTAGCGGTGGTTTTTTTGTTTGCAAGCAGCAGATTACGCGCAGAAAAAAAGGATCTCAAGAAGATCCTTTGATCTTTTCTACGGGGTCTGACGCTCAGTGGAACGAAAACTCACGTTAAGGGATTTTGGTCATGAGATTATCAAAAAGGATCTTCACCTAGATCCTTTTAAATTAAAAATGAAGTTTTAAATCAATCTAAAGTATATATGAGTAAACTTGGTCTGACAGTTACCAATGCTTAATCAGTGAGGCACCTATCTCAGCGATCTGTCTATTTCGTTCATCCATAGTTGCCTGACTCCCCGTCGTGTAGATAACTACGATACGGGAGGGCTTACCATCTGGCCCCAGTGCTGCAATGATACCGCGAGACCCACGCTCACCGGCTCCAGATTTATCAGCAATAAACCAGCCAGCCGGAAGGGCCGAGCGCAGAAGTGGTCCTGCAACTTTATCCGCCTCCATCCAGTCTATTAATTGTTGCCGGGAAGCTAGAGTAAGTAGTTCGCCAGTTAATAGTTTGCGCAACGTTGTTGCCATTGCTACAGGCATCGTGGTGTCACGCTCGTCGTTTGGTATGGCTTCATTCAGCTCCGGTTCCCAACGATCAAGGCGAGTTACATGATCCCCCATGTTGTGCAAAAAAGCGGTTAGCTCCTTCGGTCCTCCGATCGTTGTCAGAAGTAAGTTGGCCGCAGTGTTATCACTCATGGTTATGGCAGCACTGCATAATTCTCTTACTGTCATGCCATCCGTAAGATGCTTTTCTGTGACTGGTGAGTACTCAACCAAGTCATTCTGAGAATAGTGTATGCGGCGACCGAGTTGCTCTTGCCCGGCGTCAATACGGGATAATACCGCGCCACATAGCAGAACTTTAAAAGTGCTCATCATTGGAAAACGTTCTTCGGGGCGAAAACTCTCAAGGATCTTACCGCTGTTGAGATCCAGTTCGATGTAACCCACTCGTGCACCCAACTGATCTTCAGCATCTTTTACTTTCACCAGCGTTTCTGGGTGAGCAAAAACAGGAAGGCAAAATGCCGCAAAAAAGGGAATAAGGGCGACACGGAAATGTTGAATACTCATACTCTTCCTTTTTCAATATTATTGAAGCATTTATCAGGGTTATTGTCTCATGAGCGGATACATATTTGAATGTATTTAGAAAAATAAACAAATAGGGGTTCCGCGCACATTTCCCCGAAAAGTGCCACCTGACGTC

With the multiple cloning sites

...ACAAGTAATAAGCTAGCACACCGGTCGTCTAGCAATGCGATCCACGTACGTAGGATCCC

***Cerulean	NheI	AgeI	BsiWI	BamHI***

GACTGTG….

**Figure S1.** Plasmid Map of pFmiR-empty and sequence.

**pFmiR-B3GLCT-3’UTR plasmid map and 3’UTR sequence**

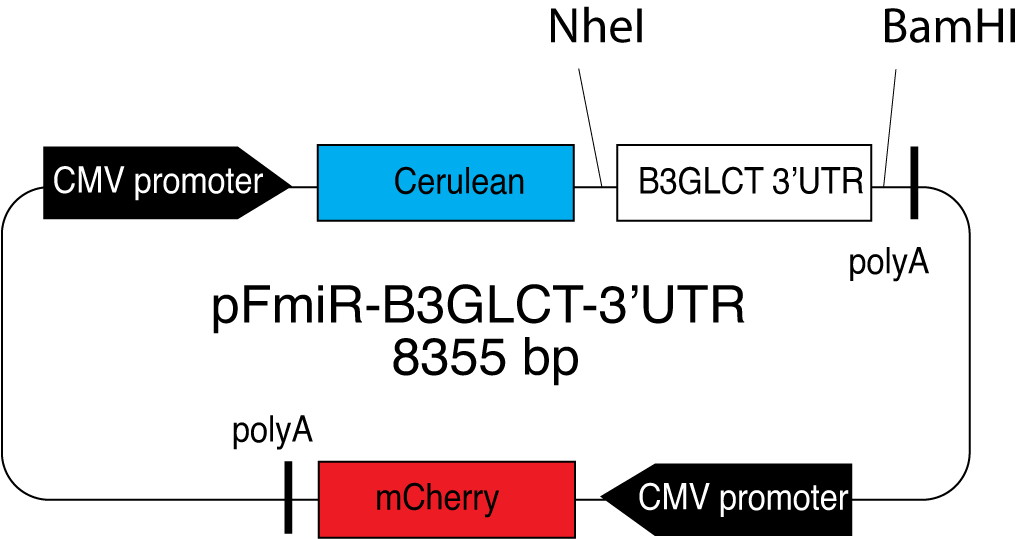

3’UTR sequence:

GCTAGCATCAGGGTGACCTGTGCGCCTAGCCTGCTCAGGGAGTGAACTGGAGACTGTGGCCTCATCCCACTGTGCTGTGCTCACAACACTTGTGTCTGCCACATGGCATTGGGTGCTTCCTGACTTTAGGGGGAGATTTTATGTATGGTATTTTTTGACAGAGGAAGAAAAGGGGTCACAGGAGAAACATTTTTTTTTCTGGGAAAAATCACTTGCTTTTGACTTATGCAGTTGTTTTAACACTTAGTGATGACTGTGTATTCTCCAAGCTGTGATACAGCAGTTTTTTTTTATTGTCACAGGGAAATAAATGGTACCAGAAGTCCCTTTCCTGTTCTGTCTCTTCATTGTAATGGAAGTTTCAGTTGGGCATGAGCCTGGAGAGATGTGACTGTCTACAGTTCTATTTGTATATATAAAAAGAAGACTGAAAGTCTTTTGACATGGATATTGTGAATGGTATGAACTTTTAAACCATATTATTGATGATGAAAATTATTTCCTGGGAACTCAGTAGGAATAATACCGTATTAAGGAATAATACTGTACATAAAACATCATGAAACCCTAGATATGAAATCCCCTGAAGTCTGTAATCATGGTGGTTATGTTTTGTCTATTCTTTTGCTGTTTGTGCCTCATAAAAAGAGAATGAGGTCTTCTGCTAGAGCTTCGTATTGCTTTGGAAGTTCATCTGTGTTTTATTTCTCCCTGAAGCCCTATCTTTATGGCTTACTTGTAACATGAAAGTAGTAGATGCTGCCAGAAAATAGTGTCCTCAATATTTTAAAACAATGTTGACATGTTTTGTTCAAGTCAGCAAGCTCTATGTGAGTCTCAGGAAGTGAATTAAATTTGGACCTTATGTTTTACTCTTGTTTTTTTTTTTTTTTTAAATGTTACTTAATGACTCTCTCCTGACTCAGGAGAGAAACCCCTTGTGGAAGGACAGCATGGTGATCAGGCAATTTCTCTGGGTTCCCAAAGAATGACATTTGAACACAGTATTTTGAAACAGCTCTAGTTTTCAAATTATATCTTTAATATATAGTAATGTAACATATTCAGTATTAATGTATAAAAAGCACTCTAATTATATAATTCAGTTTTTGTAAAGGTATTTGCATAAAATTTAATATGTCTTAAACTAATTTTGGTAAATTACTTCTTTTTTTTCTTTTTAATAAAAACTGTTACTCATTAACTTTGCTTATAATGCTTTTTATAGCCCAGCACAGAATTTAAAGCCATACCACCAAAAGTACCTGTGTGTGTTAATATGTTTTTCTTGTAGCATAGATTGACTATTTGCAATAGTATTAGTATTTACCATTTTTCCAAATTAGCAACTACCAGACCTCACGTGTTGCAGTGATAACACAATGCATTGGATTCAGTTTTGTGAAAATGGATTCTGTGGCCATCCAAGGGATGTATCAGGGATGATCAGCTGATGAGAGGCTCCAGAAGGATTTCTAGATCGCTTCAAGCCTATACTGATGGCCTTAGCTTTGTTCAGTCATTGTAACTGGGATTGTTGTCATTGCTACCGTGGTAGTCACCTTCATGTCATCTATAATAGTACTCCTGGAGAGCCCTGGCTGCCTACACCAGTGGAAAAGAGTCTCCAGTTCTGCTCTGGCCTACTAACTGTTACCACTGAGAGAACAACATGTTCATTTGACATGATTGAAGCTGGCATCCGTATATGAAGATCCTTGTCAAGCTTTCTTCTGTGGTCTGATTAGTGCCTTCTACTGATACCGGGGCACCTCCTCTGGTACTTTTAAGTGTTTTGTTAATTATATTTACTTTTTGGAATGGTGTAAGCCTAACCACAAGTAAAAGATCTTTGCCTAAGTTTTTGATTTCTCAAATATTGTGTTCATTAGTCTAGACTGGGAATGGGGAGGGGAAATGGGGAAAATGAATGAATGAAATCAGAAAAAAGTCAGCGGCTCAGTAAATACAGTTTAAAGAGAGAATAATTACTTCAGAGCTACCCTTTTAAGAGAAAACCATCAGAAATTGATAATGTTTATATAAAGTTTATAAAGCCATTGTGTTTTGTTATATAACAAATCAGAGATGTTATTTTAGAATCGATTCCCATCTAAAGAACTCAATTTTGAGTCTGACATTTCCAGGACCAGATATTGTCTTACTCACATTTCCTTTGCTTTGAAATAGGGCTTTCCTTCCAAATGGCTATTTTTAGGCTAGGGATGTTAACATCAGGGATTTGTGTGTGGAATAACTGGAATGTCATTTTTGCTTTTAAGCCATTTCTGATGATATAGCCAAAGCAGGTTGTCTGACTATGTAGGATTTTTACATCTTGAAACTAAATCAGAAATCCAGACATGAAAATAACCTTTCTAGAATGCCTAGGAGCAGAAAACAATAATAGCATGCTAAATCACAAATGATGCTATGTATGGGTATGTAAATATCAGTGCTGTCTGCATTTCTGGGTTTATTGAAGACCTCTTGTTGTATATATCCTCAAAAATTAATGTAATTGACATCTTCAAGAATGTTTCTATTGTCTTCCATTCATAATCAGAGATGTAATTTGTATGGACTAAATAAAAACTTTATTATGTAATGAAAAG

**Figure S2.** Plasmid Map of pFmiR-B3GLCT-3’UTR and sequence of 3’-UTR of B3GLCT.

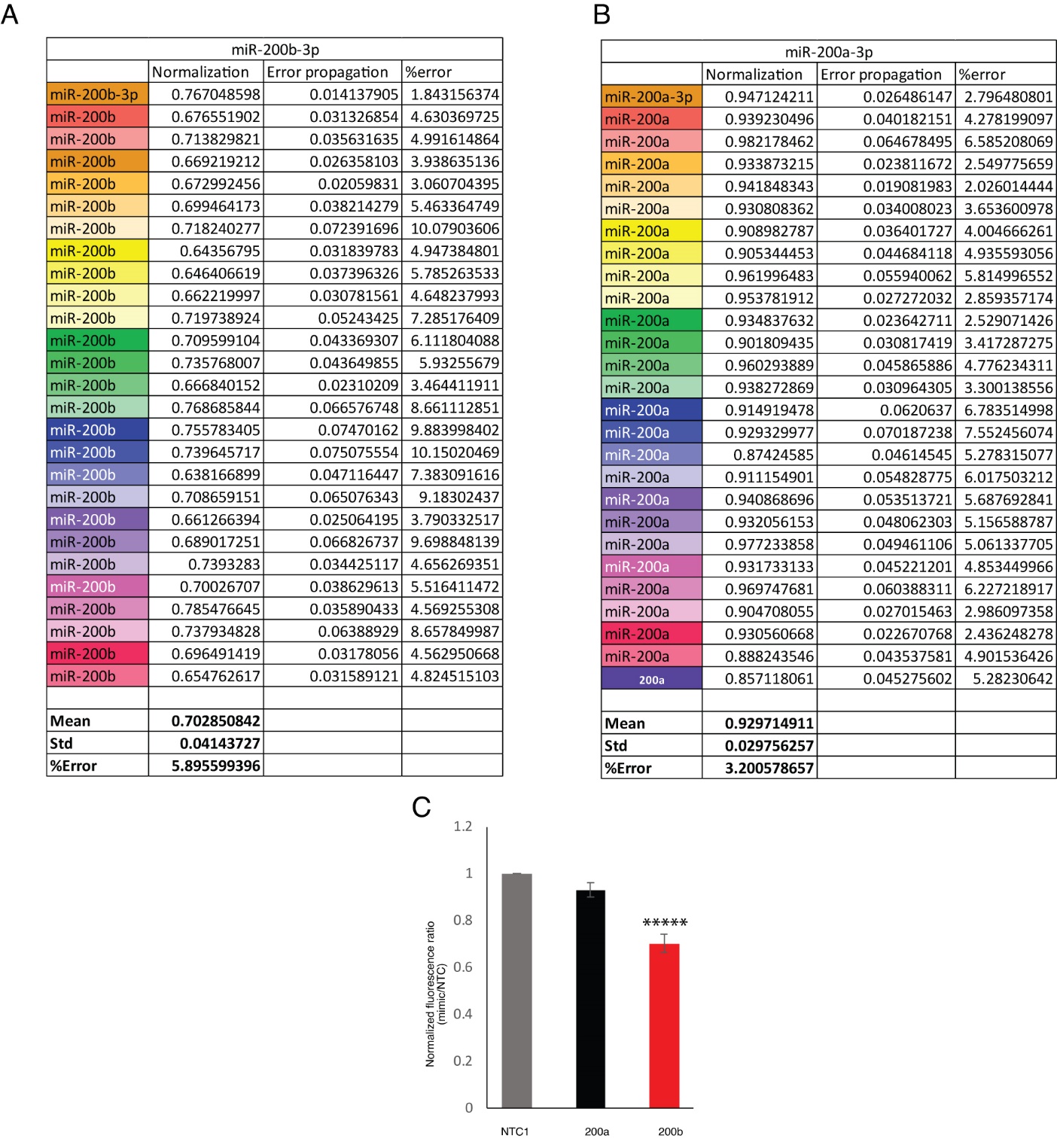

**Figure S3**. Normalization data and quantitative analysis of miR-200b-3p (A) and miR-200a-3p (B) normalized to NTC1 with triplicates for each data point over 27 384 well-plates. (C) Bar graph represented the reproducibility of the assay. Unpaired Student’s t test; *****p << 0.0001.

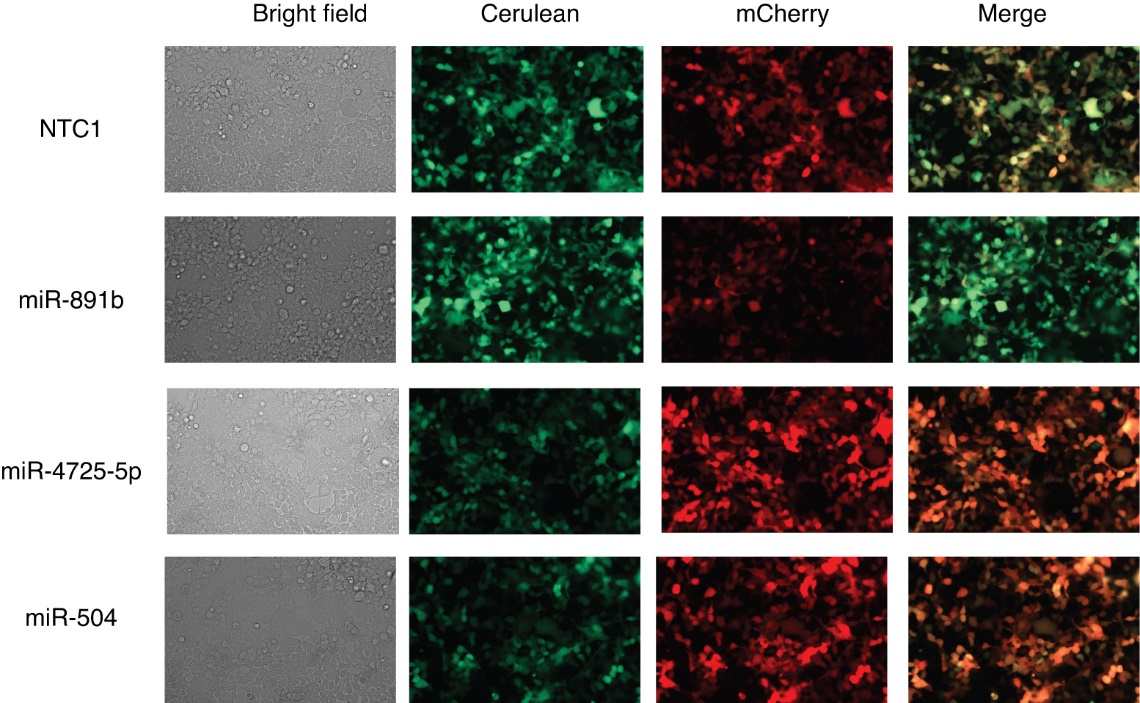

**Figure. S4**. Fluorescence microscopy images of co-transfection of pMIR-B3GLCT with NTC1, miR-891b, miR-4725-5p and miR-504 48 hours post-transfection.

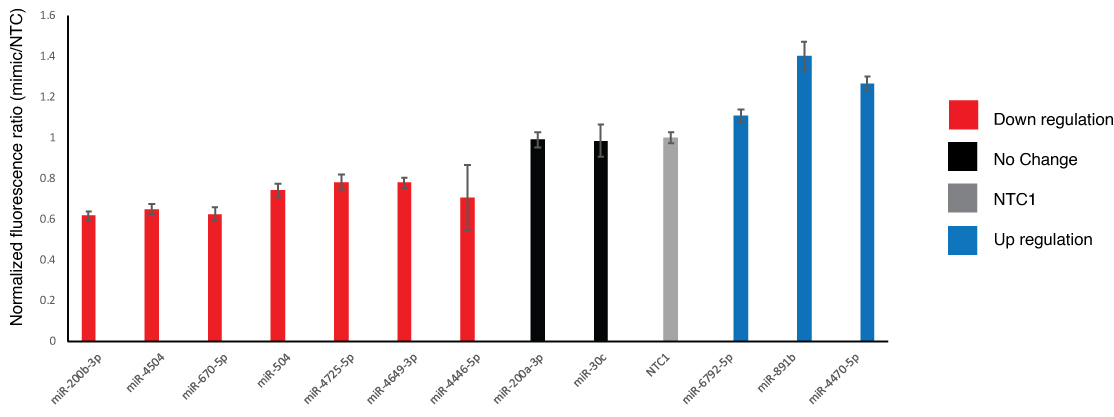

**Figure S5**. Small scale validation of miR subsets. Cells were co-transfected with pFmiR-B3GLCT and indicated miRs.

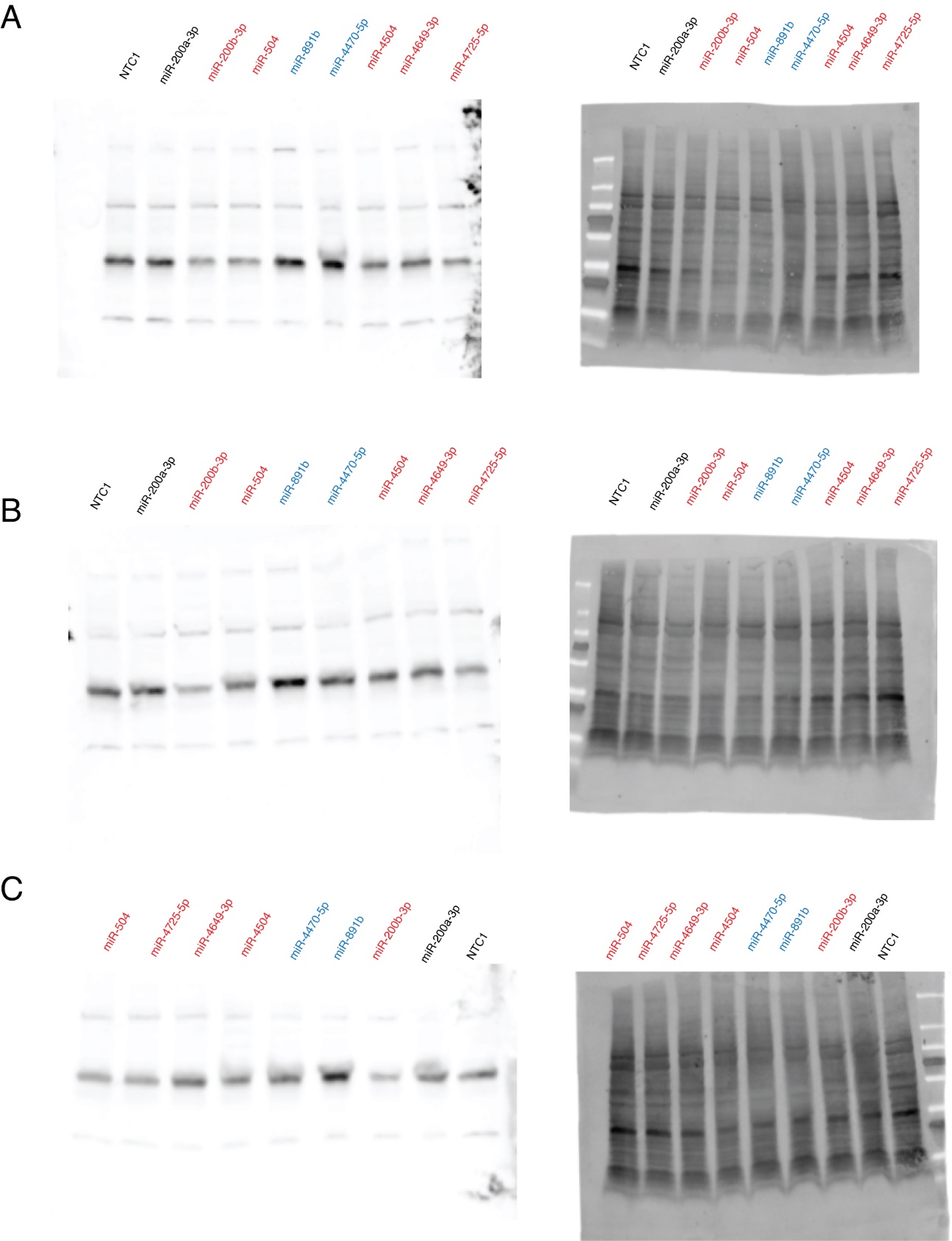

**Figure S6**. (A-C)­­ B3GLCT Western blot analysis and accompanying Ponceau S stain.

**Table S1**. Quantification of Western blot (n=3). 1,2 and 3 correspond to A, B and C in Figure S6.

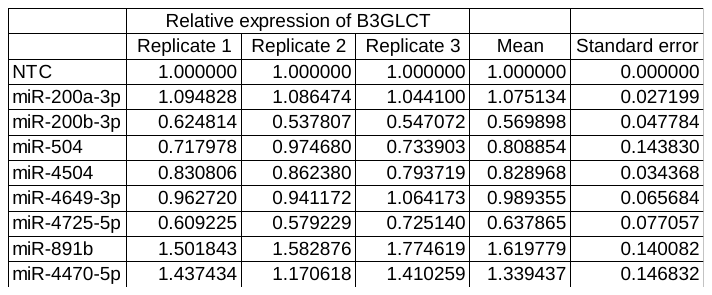

**Table S2**. Disease enrichment analysis of 27 downregulatory miRs for B3GLCT in miRNet

| Pathway | Total | Expected | Hits | Pval |
| --- | --- | --- | --- | --- |
| Preauricular dimple | 17 | 2.15 | 9 | 7.49E-05 |
| Preauricular sinus | 17 | 2.15 | 9 | 7.49E-05 |
| Malignant mesothelioma | 70 | 8.87 | 21 | 9.22E-05 |
| Adult onset | 33 | 4.18 | 13 | 9.65E-05 |
| Cleft Palate | 107 | 13.6 | 28 | 0.000101 |
| Acquired scoliosis | 153 | 19.4 | 36 | 0.000122 |
| Preauricular Fistulae, Congenital | 18 | 2.28 | 9 | 0.000133 |
| Curvature of spine | 155 | 19.6 | 36 | 0.000162 |
| Uranostaphyloschisis | 86 | 10.9 | 23 | 0.000296 |
| Frontal bossing | 82 | 10.4 | 22 | 0.00038 |
| Flatfoot | 33 | 4.18 | 12 | 0.000432 |
| Congenital small ears | 29 | 3.67 | 11 | 0.000492 |
| Hemangiosarcoma | 11 | 1.39 | 6 | 0.00106 |
| Acquired flat foot | 32 | 4.05 | 11 | 0.00128 |
| Muscle Cramp | 32 | 4.05 | 11 | 0.00128 |
| Adenoma | 19 | 2.41 | 8 | 0.00133 |
| Micrognathism | 160 | 20.3 | 34 | 0.00136 |
| Mandibular hypoplasia | 160 | 20.3 | 34 | 0.00136 |
| Hypoplastic mandible condyle | 160 | 20.3 | 34 | 0.00136 |
| Lymphoma, Follicular | 12 | 1.52 | 6 | 0.00189 |
| Intellectual Disability | 378 | 47.9 | 67 | 0.00195 |
| Sezary Syndrome | 20 | 2.53 | 8 | 0.00197 |
| Dull intelligence | 330 | 41.8 | 59 | 0.00296 |
| Orbital separation excessive | 131 | 16.6 | 28 | 0.00321 |
| Low intelligence | 326 | 41.3 | 58 | 0.00359 |
| Mental deficiency | 326 | 41.3 | 58 | 0.00359 |
| Colonic Neoplasms | 79 | 10 | 19 | 0.00371 |
| Poor school performance | 327 | 41.4 | 58 | 0.00385 |
| Ventricular Septal Defects | 63 | 7.98 | 16 | 0.00424 |
| Aplasia/Hypoplasia of the lungs | 10 | 1.27 | 5 | 0.00468 |
| Mental Retardation | 330 | 41.8 | 58 | 0.00473 |
| Small head | 192 | 24.3 | 37 | 0.00511 |
| Liver neoplasms | 70 | 8.87 | 17 | 0.00532 |
| Gastroesophageal reflux disease | 43 | 5.45 | 12 | 0.00565 |
| Global developmental delay | 313 | 39.6 | 55 | 0.00592 |
| Byzanthine arch palate | 113 | 14.3 | 24 | 0.00667 |
| Cognitive delay | 315 | 39.9 | 55 | 0.00677 |
| Squamous cell carcinoma | 66 | 8.36 | 16 | 0.00688 |
| Mental and motor retardation | 316 | 40 | 55 | 0.00724 |
| Heartburn | 39 | 4.94 | 11 | 0.00728 |
| Low posterior hairline | 34 | 4.31 | 10 | 0.00753 |
| Distal amyotrophy | 29 | 3.67 | 9 | 0.00756 |
| Bladder Neoplasm | 50 | 6.33 | 13 | 0.00774 |
| Muscle hypotonia | 311 | 39.4 | 54 | 0.00814 |
| Neoplastic Cell Transformation | 57 | 7.22 | 14 | 0.00988 |
| Precancerous Conditions | 46 | 5.83 | 12 | 0.01 |
| Renal Insufficiency | 46 | 5.83 | 12 | 0.01 |
| Alcoholic Intoxication | 16 | 2.03 | 6 | 0.0105 |
| Abnormally-shaped vertebrae | 16 | 2.03 | 6 | 0.0105 |
| Renal failure in adulthood | 41 | 5.19 | 11 | 0.0108 |
| Isolated cases | 36 | 4.56 | 10 | 0.0115 |
| Bilateral fifth finger clinodactyly | 58 | 7.35 | 14 | 0.0116 |
| Curvature of little finger | 58 | 7.35 | 14 | 0.0116 |
| Proximal muscle weakness | 26 | 3.29 | 8 | 0.0123 |
| Dilated ventricles (finding) | 65 | 8.23 | 15 | 0.0139 |
| Diffuse Large B-Cell Lymphoma | 17 | 2.15 | 6 | 0.0145 |
| Broad thumbs | 17 | 2.15 | 6 | 0.0145 |
| Cataract | 121 | 15.3 | 24 | 0.0156 |
| Liver carcinoma | 121 | 15.3 | 24 | 0.0156 |
| Downward slant of palpebral fissure | 90 | 11.4 | 19 | 0.0158 |
| Prostatic Neoplasms | 260 | 32.9 | 45 | 0.0159 |
